# An intracortical brain-machine interface based on macaque ventral premotor activity

**DOI:** 10.1101/2025.06.11.659174

**Authors:** Sofie De Schrijver, Jesus Garcia Ramirez, Santiago Iregui, Erwin Aertbeliën, Joris De Schutter, Tom Theys, Thomas Decramer, Peter Janssen

## Abstract

The majority of brain-machine interface (BMI) studies have focused on decoding intended movements based on neural activity of primary motor (M1) and dorsal premotor cortex (PMd). The ventral premotor cortex (PMv), and more specifically area F5c, has been implicated in object grasping and action observation, and may represent an alternative for motor BMI control due to its phasic modulation during action observation. Using chronically implanted Utah arrays in F5c, PMd, and M1 in two male macaques, we compared the efficacy of controlling a motor BMI based on neural activity of each area. PMv decoding reached similar or even higher success rates than M1 and PMd in a 2D cursor control task, especially when controlling for the number of motion selective channels that were used by the decoder. We found similar results during a 2D robot avatar control task in a simulated 3D environment. At both the multi-unit and the population level, neural responses were highly similar during the training phase (passive observation of cursor movements) and the online decoding phase, and only a small subset of neurons modulated its selectivity for the direction of motion. Thus, ventral premotor area F5c may represent an alternative for online motor BMI control.

**Significance statement:** We present the first study on online cursor and robot avatar control using neural activity of ventral premotor cortical area F5c. Known for decades for the presence of mirror neurons, which are active during both action execution and action observation, area F5c can support online BMI control with performance comparable to that of dorsal premotor and primary motor cortex. The population dynamics in all three areas were highly similar between the training phase and the online decoding phase.

## Introduction

Intracortical brain-machine interfaces (BMIs) for motor control have been remarkably successful in the past two decades. Decoding motor cortical activity in nonhuman primates and humans has led to online 3D cursor control (Taylor, Tillery, and Schwartz 2002; Jarosiewicz et al. 2008), robot control for self-feeding (Velliste et al. 2008; Hochberg et al. 2013), movement of paralyzed muscles by means of electrical stimulation (Ajiboye et al. 2017), locomotion through a brain-spine interface (Lorach et al. 2023), and even hand writing and speech (Willett et al. 2021; 2023). The overwhelming majority of these studies (except (Sakellaridi et al. 2019)) have recorded neuronal activity in primary motor cortex (M1) and to a lesser extent dorsal premotor cortex (PMd).

In cases where M1 itself is damaged (e.g. after a stroke, or in patients with Amyotrophic Lateral Sclerosis), it is important to be able to use activity in other cortical areas for online BMI motor control. A previously unexplored candidate area for online cursor control is ventral premotor cortex (PMv), more specifically its F5c subsector that is located on the frontal convexity. The macaque F5c contains many neurons that are active during grasping movements and during action observation (di Pellegrino et al. 1992; Gallese et al. 1996), reversible inactivation of this area causes motor slowing (Fogassi et al. 2001) and some F5c neurons encode different types of grasp (e.g. precision grip vs power grip (Raos et al. 2006). An additional advantage of using a premotor area rather than M1 might be that it could be easier for the subject to control a cursor or a robot based on cortical activity reflecting high-level motor planning commands (see also (Aflalo et al. 2015)).

We recently studied Action Observation/Execution neurons (AOENs) in F5c with videos of grasping actions and reduced control videos (De Schrijver, Decramer, and Janssen 2024), and found that the large majority of these AOENs did not respond throughout the action video but rather fired in a phasic way in specific epochs of the action video. Overall, these neurons seemed to signal the position of the hand (or that of an abstract ellipse stimulus) with respect to the to-be-grasped object, which could be very useful for online cursor control. Although the potential of AOEN activity for BMI applications has been suggested (Tkach, Reimer, and Hatsopoulos 2007), no study has investigated the performance of a BMI based on F5c activity for cursor or robot control. Here, we explored the potential of F5c activity for a motor BMI and compared its decoding performance to neighboring areas PMd and M1. We implanted a 96-electrode Utah array in each of these areas and trained decoders with the activity recorded in each of these areas during the active execution of a center-out task and during passive fixation of cursor movements on a display. In general, we achieved accurate online cursor control using each of the three areas. Remarkably, both the success rate and speed of F5c decoding reached a similar level as that based on M1 activity in the different tasks. Thus, F5c represents an interesting alternative cortical area for online BMI control.

## Methods

### Surgeries and electrophysiology

Two male rhesus monkeys (*Macaca Mulatta*) were chronically implanted with three 96- channel microelectrode Utah arrays (Figure 1A) with an electrode length of 1.5mm and an electrode spacing of 400µm (4x4mm; Blackrock Neurotech, UT, USA). The arrays were implanted guided by stereotactic coordinates and anatomical landmarks in the hemisphere contralateral to the monkeys’ working hand. We used a pneumatic inserter (Blackrock Neurotech) with a pressure of 1.034 bar and an implantation depth of 1 mm. Postoperative anatomical scans (Siemens 3T scanner, 0.6mm resolution) verified the position of the Utah arrays in ventral premotor area F5c, dorsal premotor area F2, and the primary motor cortex (M1). Electrical microstimulation on M1 electrodes evoked movements of the contralateral hand, confirming implantation in the hand region of M1. All surgical and experimental procedures were approved by the ethical committee on animal experiments of the KU Leuven and performed according to the *National Institute of Health’s Guide for the Care and Use of Laboratory Animals* and the EU Directive 2010/63/EU.

**Figure 1:**
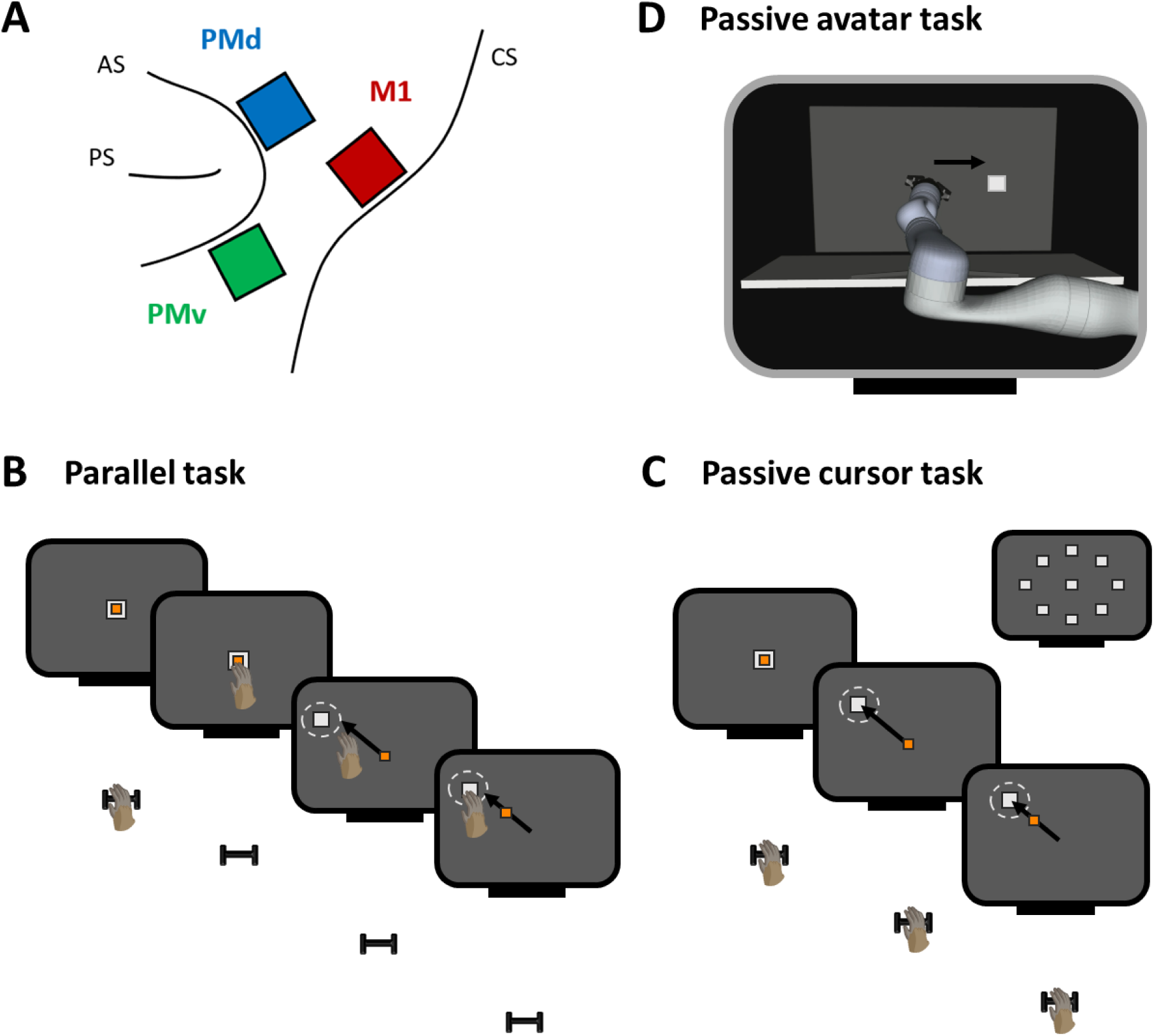
Implantation sites and experimental set-up. (A) Schematic illustration of the chronically implanted Utah arrays in ventral premotor area F5c (green), dorsal premotor area F2 (blue), and the primary motor cortex M1 (red). The indicated sulci are the principal sulcus (PS), arcuate sulcus (AS), and central sulcus (CS). (B) Schematic temporal sequence of the Parallel task. The monkey performed a center-out task while a cursor moved from the center target to the outer target. The cursor started moving after the monkey touched the center target and the center target disappeared. White square = target, orange square = cursor, white dotted line = target window. (C) Same as (B) but for the Passive cursor task in which the monkey did not move and only the cursor moved from the center target to the outer target. The inset shows the central target and eight possible outer targets. (D) Schematic illustration of the Passive avatar task in which the cursor was replaced by the virtual Kinova robot arm. The avatar arm moved from the center target to the outer target. The black arrows indicate cursor/avatar movement.

Neural activity was recorded with either a 192-channel or a 256-channel Cerebus data acquisition system (Blackrock Neurotech, UT, USA). In each recording session, we connected two arrays fully (i.e. all 96 electrodes) of which we used the signal of one array for training of the decoder. Additionally, 64 electrodes from the third array were connected during recordings with the 256- channel Cerebus data acquisition system. The signal was high pass filtered (750Hz) and sampled at 30kHz. In each recording session, the threshold to detect neural activity was manually set below the noise level for each channel to capture the activity of individual neurons. For this study, we used the online recorded activity without spike sorting, thus the activity recorded with one electrode could contain the activity of one or multiple single neurons.

### Experimental design

First, the monkeys were trained to perform a center-out task on a touchscreen (17.3 inch) that was placed in front of the monkey at a distance of 28cm, while they were head-fixed and seated in the dark. The ipsilateral arm was gently restrained. To start a trial, the monkey placed its contralateral hand on a resting position, which induced the center target (white square) to appear in the center of the screen. After touching the center target, the center target disappeared and one of eight predefined outer targets (white squares) appeared pseudorandomly. The monkey was required to touch the outer target within 1500ms. Successful trials were rewarded with a liquid reward. When the monkeys were proficient in the center-out task, we introduced the three tasks used in this study. In the first task, the monkeys performed the center-out task but a cursor (small orange square) was added (Parallel task, Figure 1B). When the monkey touched the center target, the center target disappeared and the cursor started moving towards the outer target. Thus, both the monkey hand and the cursor moved towards the outer target in the same time period, and the monkey was rewarded when both reached the target within the predefined period. Importantly, the monkey did not track the cursor movements with its hand, instead the hand movements were consistently faster, reaching the outer target before the cursor reached it. In the second task, the monkey passively observed the cursor moving from the center to the outer target and a reward was given when the cursor reached the outer target (Passive cursor task, Figure 1C). After a successful trial, the monkey was rewarded and the cursor reappeared at the center of the screen. In the third task, the cursor was replaced by a 3D-rendered avatar arm (Kinova Gen 3 robot arm) that moved from the center target to the outer target on a virtual screen (Passive avatar task, Figure 1D). The avatar was always positioned on the right side of the screen, similar to the monkey’s contralateral hand. When the avatar reached the outer target, a reward was given and the avatar moved back to its initial center position. During all tasks, eye movements were monitored using an infrared-based camera system (Eyelink 500; SR Research, Ontario, Canada) to ensure that the monkey was looking at the screen during the cursor and avatar movements.

Each task consisted of two phases: a training phase in which the cursor/avatar movements were computer-driven (Observation), and an online decoding phase in which the movements were driven by brain activity (BMI control, Figure 2A). In each recording session, we obtained approximately 200 trials (25 repetitions per target) during the training phase (i.e. when the monkey passively observed the cursor/avatar movements). Note that during the Parallel task, the monkey observed the cursor movement while also moving his hand. We then used the obtained neural data of one brain area to train the BMI decoder (alternating brain areas every day to account for learning effects across multiple sessions). Next, we ran the same task as during the training phase, but now the movements of the cursor/avatar were controlled by the monkey’s neural activity (decoding phase). The decoder predicted cursor/avatar velocities in 50ms timesteps to ensure a smooth trajectory. For all tasks, a trial was considered successful during the decoding phase when the cursor/avatar reached the target and stayed within the target window for 500ms (white dotted circles around the outer target in Figure 1B and 1C). The target window had a radius of 2.75cm in the Passive cursor/Parallel task, and a radius of 6cm in the virtual environment of the Passive avatar task. The predefined period to reach the outer target was 4 and 7.5s for the Passive cursor/Parallel task and the Passive avatar task, respectively. We implemented a longer period for the Passive Avatar task to account for the slower movement speed of the avatar.

**Figure 2:**
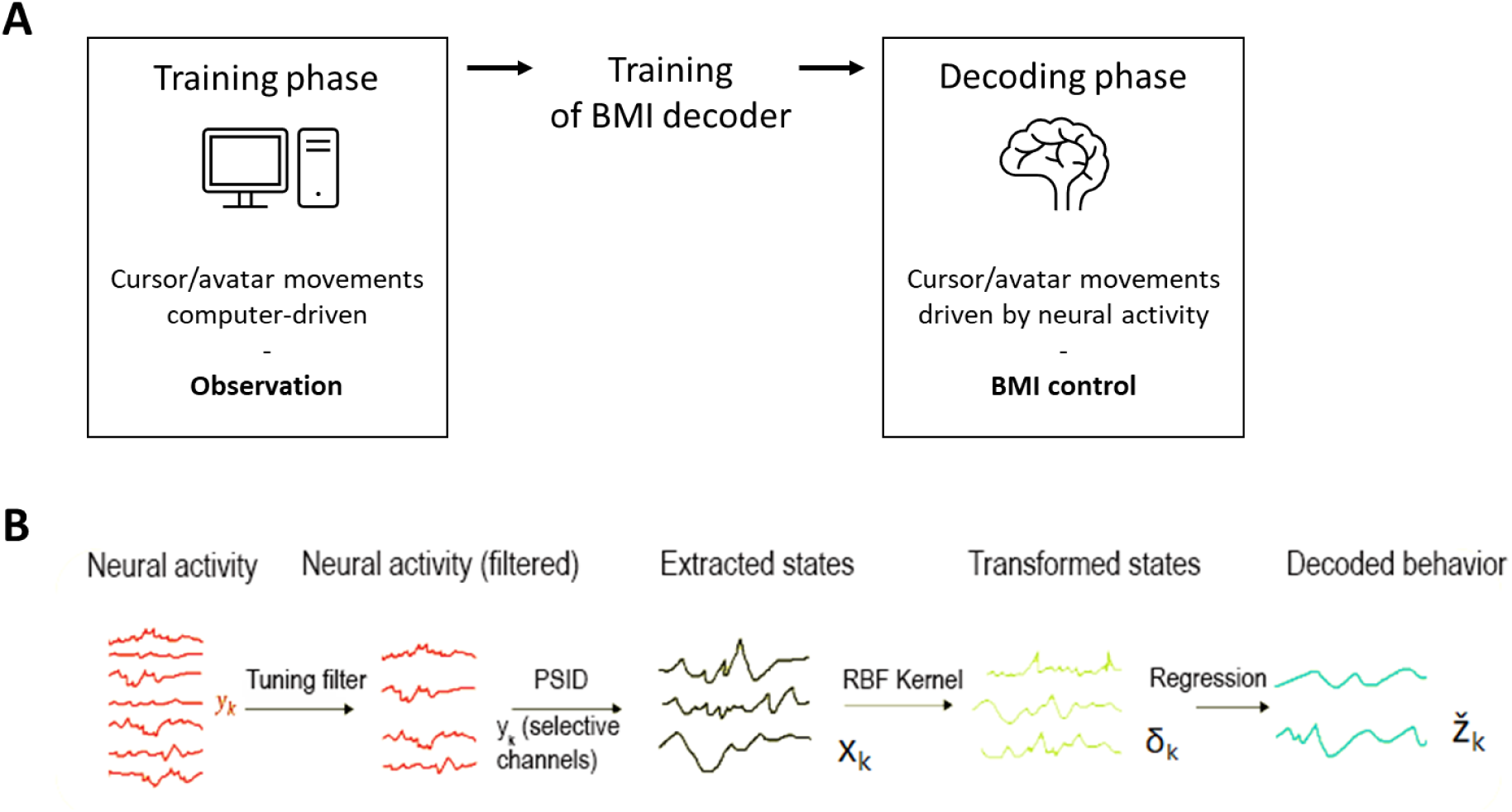
BMI paradigm. (A) Each task consisted of two phases. First, neural data was obtained during observation of the cursor/avatar movements that were computer-driven (‘Training phase’). After finishing the training phase, we trained a BMI decoder with the obtained data. Finally, the cursor/avatar movements were controlled by the BMI decoder based on real-time neural activity (‘Decoding phase’). (B) The neural activity of motion selective channels was extracted and fed into the Preferential Subspace Identification (PSID) algorithm. The extracted states were transformed with a Radial Basis Function (RBF) kernel. A regressor predicted the intended behavior, i.e. the velocities of the cursor or avatar.

For the third task, we used two variations during the online decoding phase, one in which online avatar control solely depended on brain activity (Passive avatar task) and one where online BMI control was assisted by an algorithm (Reactive Virtual Guidance Fixture, RVGF (Iregui, De Schutter, and Aertbelien 2021)). The RVGF is a shared control assistive strategy that implements a force field in the space surrounding a tube starting at the end effector and directed towards the target location. The impedance field around the tube-shaped volume acted like a variable stiffness spring. As the end effector moved away from the center of the tube, it experienced an increasing force pulling it back towards the interior. When the end effector was inside the tube, it could move freely. Note that the RVGF used position restraints, therefore allowing the avatar to also move outside the tube (Passive assisted avatar task). We implemented the RVGF to improve decoder performance in the second variation of the Passive avatar task, since we observed low performance when avatar control relied solely on brain activity.

### Neural decoder

#### Decoding Pipeline

Our decoding pipeline leveraged a non-linear extension of the Preferential Subspace Identification (PSID) algorithm (Sani et al. 2021) to translate recorded neural activity into real-time predictions of cursor/avatar velocities (Figure 2B). As a pre-processing step, threshold-crossing rates were binned in 50ms and concatenated into a 96 x 1 vector (𝑦_𝑘_). To improve signal quality and reduce computational complexity, we selected the K most motion selective channels, by calculating the median absolute Pearson correlation coefficient between its firing rate and the binned x and y velocities of the cursor/avatar along each axis. The threshold for selecting a channel γ was optimized using the first recording sessions and finally set at 0.3 for all online decoding sessions. Finally, this K x 1 vector (𝑦^_𝑘_) was fed into the PSID algorithm.

We employed the standard Python implementation of the PSID algorithm provided by Sani et al. (Sani et al. 2021). Briefly, the PSID algorithm assumes that the brain’s state at any time is a high-dimensional hidden variable that influences both the observed neural activity and the measured behavior (Figure 2B). PSID is a general dynamic linear state space model to describe the neural activity (𝑦_𝑘_ ∈ R^𝑛𝑦^) and behavior (𝑧_𝑘_ ∈ R^𝑛𝑧^) as follows:

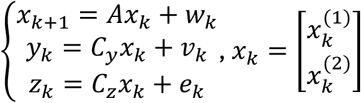

Where 𝑥_k_ ∈ R^𝑛𝑥^ is the hidden state that drives the observed neural activity, 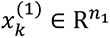 and 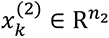 are its components that are relevant and irrelevant to behavior, respectively. Lastly, 𝑒_𝑘_ represents the behavior dynamics that are not captured by the observed neural activity, and 𝑤_𝑘_ and 𝑣_𝑘_ are noise variables. A, 𝐶_𝑦_ and 𝐶_𝑧_ are noise statistics and the model parameters, which are learned in the training phase combining neural and behavioral data. Finally, the full model separates the hidden states into 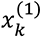 and 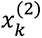 to enable PSID to focus on learning the neural dynamics that are relevant to behavior (that is, only 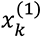). In our implementation, we only use 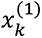 for further decoding pipeline steps.

While PSID offers a linear model, we observed improved performance by incorporating a non-linear transformation of the hidden states. This is likely because neural representations often exhibit non-linearities (Koch and Segev 2000). We opted for a radial basis function (RBF) kernel to capture these non-linear relationships. To enable real-time inference, the RBF kernel was approximated efficiently using the Nyström method (Williams and Seeger 2001) implemented in scikit-learn. This method constructs a reduced representation of the kernel matrix by subsampling data points, allowing for faster computation during online decoding. We can construct the eigendecomposition of the kernel matrix K, based on the features of the data, and then split it into sampled and unsampled data points:

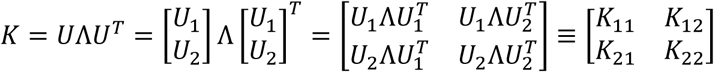

Where 𝑈 is orthonormal, Λ is a diagonal matrix of eigenvalues, 𝑈_1_ is orthonormal matrix of samples that were chosen and 𝑈_2_ is orthonormal matrix of samples that were not chosen. Finally, to map the non-linear hidden states (after applying the RBF kernel) to predicted velocities, we employed separate regularized ridge regression models for each velocity component (x and y). For each model, the features were constructed by concatenating the hidden states across all time points within a trial. The corresponding behavioral data (velocity) served as the regressor.

### Decoding Training Overview

Each day, we trained a new decoder offline using the neural data collected during cursor/avatar movement in the training phase. Based on a supervised learning paradigm, the offline training provided the decoder with an initial understanding of the relationship between neural activity and intended movement. To enhance the ability of the decoder to generalize and prevent overfitting, we generated realistic offline trajectories using Bézier curves that were used for the computer-driven cursor/avatar movements in each tasks’ training phase. These curves allowed to simulate the diverse and smooth movement patterns the decoder would need to adapt to during real-time online use.

Early in our development process, we employed a Bayesian optimization framework to systematically explore the hyperparameter space. This approach efficiently searches for optimal parameter combinations by building a probabilistic model of the objective function and using it to select promising values to evaluate. We optimized for the cross-validated offline R-squared value between predicted and actual trajectories. After establishing a set of hyperparameters that consistently yielded high performance across initial sessions, we fixed these parameters for subsequent recordings. This decision was motivated by our primary goal of enabling fair comparisons of motor BMI performance across recordings from different brain areas, rather than maximizing decoding performance within each individual session. A summary of the hyperparameters used to train our decoding pipeline can be found in Supplementary Table 1.

To address potential neural signal changes (e.g., changes in firing rates or tuning) that can occur during online BMI use, we implemented an online retraining procedure. This adaptive strategy periodically recalibrated the decoder as the monkey actively controlled the cursor or avatar, ensuring that the BMI remained accurate and responsive. We triggered the retraining process after a predetermined number of completed trials (sixteen for the Parallel and Passive cursor task, and eight for the Passive avatar task). Crucially, this process ran in the background during task performance to minimize disruption to the monkey’s control, and the newly retrained decoder seamlessly replaced the previous one. Our retraining approach employed the ReFit method (Gilja et al. 2012). We retrained the decoder with an “intention-based” kinematic training set. This approach aimed to better align the decoder with the monkey’s intended movement goals, even if the initial BMI output was not perfectly accurate. In this set, the decoder was trained using target-adjusted velocities, where predicted velocities in each trial were rotated towards the actual target direction to help the decoder learn from potential mismatches between intention and initial execution. To ensure a fair comparison between retrained decoders, the online retraining dataset maintained a fixed size of 200 trials, combining the most recent offline trials with newly acquired online trials in a first-in-first-out (FIFO) scheme. This allowed the decoder to adapt to recent neural patterns while retaining foundational knowledge from the offline training. Note that the number of motion selective channels could change during the online retraining.

## Statistical analysis

All data were analyzed using custom written Matlab (the MathWorks, R2019b) and Python scripts (Python 3.12). The success rate was calculated for each recording session as the number of correct trials divided by the total number of trials that were not aborted (i.e. trials stopped before completion because the monkey stopped fixating). To account for variability in the quality of the neural data and thus in the daily performance, we calculated the running average of the success rate over time by taking the sum of the success rate of three consecutive sessions with a sliding window of one session. The time-to-target of each trial was defined as the time for the cursor to reach the target and stay within the target window for 500ms. This value could not exceed the maximal predefined window of 4s. To control for the number of motion selective channels in the different areas over time, we calculated the success index by dividing the success rate by the number of selected channels for each session. To assess the difference of success indices between the motor areas, Kruskal-Wallis tests with post-hoc pair-wise comparisons with Dunn-Sidak correction were used.

For each trial, we calculated the raw spike rate from 200ms before the start of the cursor movement until the end of the cursor movement. Since the cursor movements differed in length in both the training phase (due to the curved trajectories) and the online decoding phase, time was defined in arbitrary units. Neural activity during the cursor movement was interpolated to 60 units for visualization purposes. To investigate the direction selectivity in the recorded areas, we determined the d prime (d’) index for the activity of each electrode.

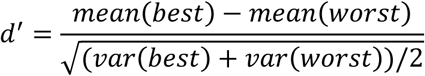

With best = the average activity during cursor movement towards the target with the highest overall response, and worst = the average activity during cursor movement towards the target with the lowest overall response. We calculated the d’ indices during both the training phase and the decoding phase to assess whether sites changed their selectivity for the direction of motion during passive observation and active BMI control. To compare the change in d’ between the areas, we used the two-sample Kolmogorov-Smirnov test to compare the distributions of the d’ changes. The d’ change was calculated for each site by subtracting the d’ of the training phase from the d’ of the online decoding phase. Thus, positive values indicate an increase selectivity for direction of motion during active cursor control compared to passively observing the cursor.

To assess the degree of alignment between the neural activity in the training phase and the online decoding phase, we performed a subspace overlap analysis (Elsayed et al. 2016). For each recording session and each area, we constructed matrices T and D with the neural activity for the training and the decoding phase respectively, that were both n x t, with n the number of electrodes that recorded significant task-related activity in both phases and t the number of time points. Each row of the matrices contained the normalized (z-scored) average neural activity. When an area contained less than 10 electrodes with responsive activity in a recording session, the data were excluded from the analysis. Next, we performed principal component (PC) analysis on matrix T and D to obtain the first ten training PCs and the first ten decoding PCs, respectively. We then projected the training activity (T) and the decoding activity (D) onto the training PCs to visualize the neural trajectories in a shared 3D neural space. The trajectories were smoothed by taking the running average with a window of three and a sliding window of one. To quantify the amount of shared variance between the neural activity of the training and the decoding phase, we calculated the sum of the variance captured when projecting T onto each of the first ten decoding PCs and the sum of the variance captured when projecting T onto each of the first ten training PCs. The alignment index (AI) was calculated by taking the ratio of both sums. Thus, a low AI value indicates that the training and decoding subspaces are approximately orthogonal, as only limited variance will be captured by projecting T onto the decoding PCs. AI values range from 0 (perfect orthogonality) to 1 (perfect alignment).

## Results

To investigate the potential of macaque ventral premotor area F5c in controlling a motor BMI, and to compare its performance to that of M1 and PMd, we implemented three tasks (Figure 1B-D). In the first task, the monkey executed a center-out task while controlling cursor movements with a BMI decoder trained on neural data of one of the three areas (Parallel task). In the second task, the hand remained still and only the cursor needed to be moved (Passive cursor task). In the third task, the cursor was replaced by an robot avatar arm in a virtual environment and the monkey controlled the avatar movements. Online decoding in this task either depended entirely on brain activity (Passive avatar task) or was assisted by an assistive algorithm (Assisted Passive avatar task). Cursor/avatar velocities were predicted online in 50ms timesteps. Every recording session used the signal from only one of the areas to explore interareal differences over sessions. Monkey V. performed 42, 33, 39, and 26 sessions of the Parallel, Passive cursor, Passive avatar, and Passive assisted avatar task, respectively with on average 12 sessions per area (range [9 16]). Monkey L. performed 48, 39, 21, and 21 sessions of the Parallel, Passive, Avatar, and Assisted Passive avatar task, respectively with on average 11 sessions per area (range [7 18], Supplementary Table 2).

## Online performance during the Parallel and Passive cursor task

Figure 3A shows cursor trajectories of 16 trials from example sessions of the Parallel task recorded in each of the three areas (two trials per target, monkey L.). The circles indicate the electronically defined target windows and the dots inside each circle indicate the target locations. Figure 3B shows the improvement of the decoder performance within a recording session (from PMd) due to the online retraining. These cursor trajectories were obtained at the beginning and towards the end of the recording session after multiple instances of retraining the decoder. Initially, online decoding performance was at chance level (first sixteen trials, left panel in Figure 3B), but rapidly improved after seven retraining instances (last sixteen trials, right panel in Figure 3B).

**Figure 3:**
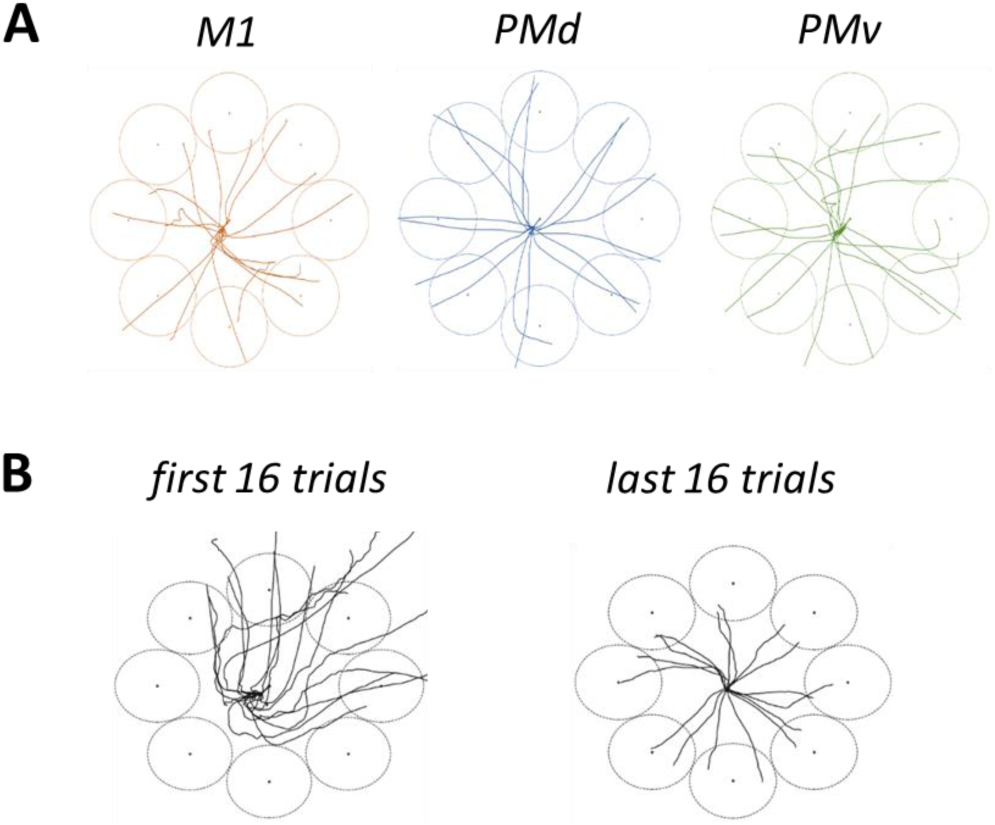
Cursor trajectories during online decoding of the Parallel task. (A) Cursor trajectories of 16 trials from the center target to the eight possible outer targets (dot) with their respective target window (circle). Each example shows trials from a recording session that used the indicated area for online decoding. (B) Cursor trajectories of the first 16 trials and the last 16 trials in an example recording session in PMd.

To ensure that long-term training effects or changes in signal quality did not affect the results, we alternated between each of the three areas on consecutive days and plotted a running mean of the success rate as a function of session number for the two monkeys separately (Figure 4). In the Parallel task, monkey V. already reached an average success rate above 55% in the first three sessions when using M1 and PMd activity, but performed much less when using PMv (χ^2^(1,678) = 55.79, p<0.001 relative to M1, and χ^2^(1,761) = 191.81, p<0.001 relative to PMd; Figure 4A, left panel). However, PMv-based decoder performance improved over the next sessions and even reached a higher level than M1 in the last part of the recording sessions (χ^2^(1,1304) = 6.72, p = 0.0095). In the second half of the recording period, the decoder performance of PMd decreased whereas both M1 and PMv continued to achieve above 50% correct responses. In the Passive cursor task of monkey V. (right panel in Figure 4A), the differences between the three areas were less pronounced in the first half of the recording period, but PMv improved remarkably and even outperformed M1 and PMd in the last five recording sessions (χ^2^(1,1193) = 129.77, p<0.001 relative to M1, and χ^2^(1,1066) = 120.95, p<0.001 relative to PMd). In monkey L., the overall online success rate in the Parallel task was higher, especially for M1 and PMd in the first recording sessions, but again PMv decoding performance reached a level similar to M1 and slightly lower than PMd in the second half of the recording period (χ^2^(1,1299) = 0.20, p = 0.6516 relative to M1, and χ^2^(1,1845) = 59.52, p<0.001 relative to PMd). The changes in performance were even more striking in the Passive cursor task, where we initially measured a very low PMv-based decoder performance and a significant improvement (χ^2^(1,1450) = 89.20, p<0.001) reaching the level of M1 in the second half of the recording period (χ^2^(1,1299) = 0.20, p = 0.6516). In monkey L., PMd-based decoder success rate remained high throughout the recording period in both tasks. Thus, accurate online cursor control based on neural activity recorded in PMv is possible and can reach a similar level as M1 and in some cases even PMd.

**Figure 4:**
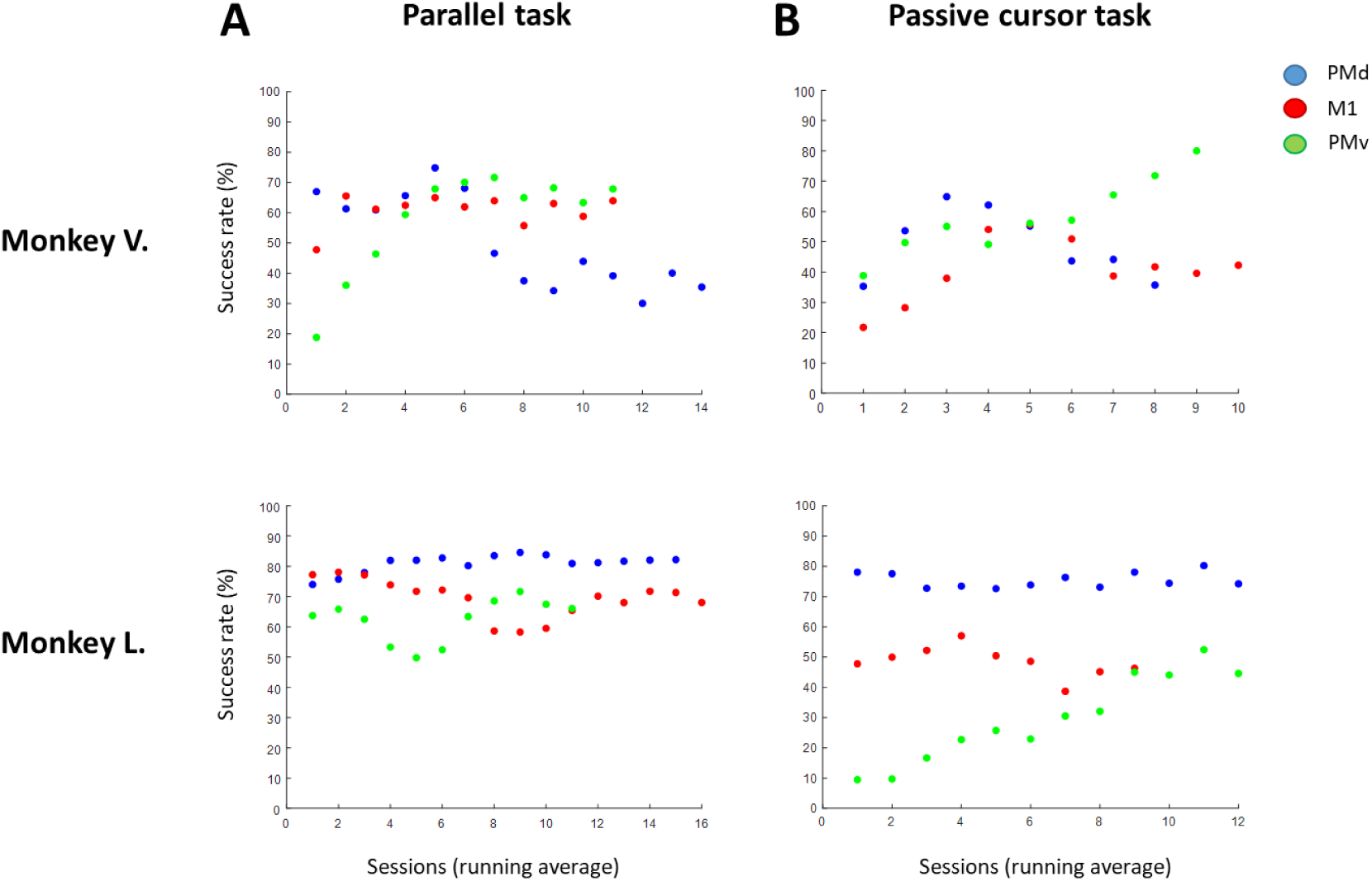
Performance in the Parallel and Passive cursor task during online decoding. (A) Running average of the success rate over time (average of three sessions with a sliding window of one session) for each monkey in the Parallel Task. (B) same as (A) but for the Passive cursor task. Blue = PMd, red = M1, and green = PMv.

We also investigated whether the three areas differed in the time necessary for the cursor to reach the target during online decoding. In both monkeys and for all three areas, the time-to-target was significantly shorter in the Parallel task compared to the Passive cursor task (mean difference ranging from 262 to 663ms, Mann-Whitney U tests, all p-values < 0.05). However, the mean time-to-target did not significantly differ between the three areas in Monkey V. (Mann-Whitney U tests, all p-values > 0.05). Monkey L. was significantly faster when using PMd activity although the difference with the other areas was modest (189ms compared to M1 and 244ms compared to PMv for the Passive cursor task, and 46ms compared to M1 and 74ms compared to PMv).

### Online performance during the Passive avatar task

Moving a cursor towards a target may not be an optimal task to recruit neurons in PMv, where Action Observation/Execution neurons (AOENs) have been discovered responding during object grasping and during passive observation of a video showing a hand grasping an object (Caggiano et al. 2011). Therefore, we tested whether online decoding performance based on PMv activity would improve in a more realistic task in which an robot avatar arm had to be moved towards a target. The overall performance in this task was markedly lower compared to the parallel and passive cursor task (Figure 5A and B, left panels). In monkey V., online decoding based on PMv activity reached a higher performance as PMd (χ^2^(1,1475) = 8.74, p = 0.003 relative to PMd) but a slightly lower performance as M1 over all recording sessions (χ^2^(1,1858) = 11.05, p<0.001 relative to M1). In monkey L., however, PMv-based decoder performance was significantly lower than that of the other two areas (χ^2^(1,1447) = 8.14, p = 0.004 relative to M1, and (χ^2^(1,1451) = 92.51, p<0.001 relative to PMd). To facilitate the online control of the robot avatar arm, we implemented an assistive algorithm (Reactive Virtual Guidance Fixture, RVGF) that guided the avatar towards the target within certain boundaries (Iregui, De Schutter, and Aertbelien 2021). The addition of this assistive algorithm improved online decoding performance in all three areas, but not above the level observed in the parallel and passive cursor tasks. Monkey V. achieved a performance level with PMv activity (near 60% success rate) that was equally high compared to M1 activity (χ^2^(1,2842) = 0.31, p = 0.5766) but significantly higher compared to PMd activity (χ^2^(1,2678) = 48.72, p<0.001), whereas in monkey L. PMv-based decoder performance remained lower than M1- and PMd-based performance ( χ^2^(1,1542) = 33.12, p<0.001 relative to M1, and (χ^2^(1,1556) = 332.65, p<0.001 relative to PMd).

**Figure 5:**
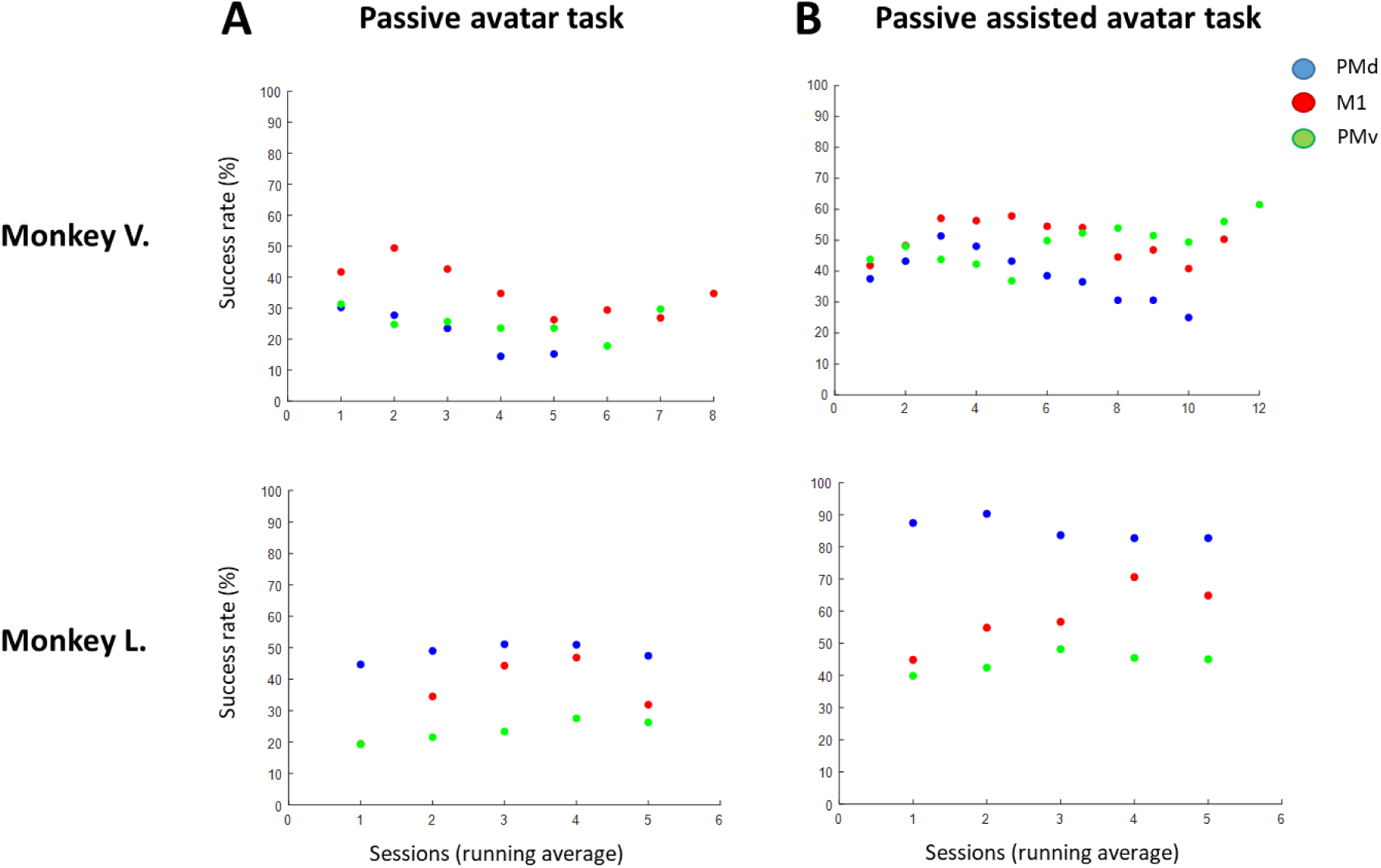
Performance in the Avatar and Passive assisted avatar task during online decoding. (A) Running average of the success rate over time (average of three sessions with a sliding window of one session) for each monkey in the Passive avatar task. (B) same as (A) but for the Passive assisted avatar task. Blue = PMd, red = M1, and green = PMv.

### The effect of the number of motion selective electrodes on online decoding

A critical factor driving decoding performance is the number of electrodes in an array used for decoding (Carmena et al. 2003). These electrodes correspond to the motion selective electrodes (K selected electrodes that meet the kinematics selectivity threshold, see methods). Although all three arrays were identical and implanted at the same time, the spike yield clearly differed between the arrays and also varied between sessions and over time. Therefore, the number of motion selective electrodes at the start of each online decoding phase differed between areas, sessions, and monkeys. On average, 33 (range [13 50]), 49 (range [30 68]), and 23 (range [10 50]) electrodes were selected for PMv, M1, and PMd, respectively, in monkey V. In monkey L., on average 29 (range [7 83]), 52 (range [14 88]), and 45 (range [33 65]) electrodes were selected for PMv, M1, and PMd, respectively. We observed that in several sessions, PMv reached a robust performance despite a relatively low number of motion selective channels. To capture this effect, we calculated the success index for each recording session as the percent correct divided by the number of motion selective channels in each task (Figure 6). In the Parallel and Passive cursor tasks, the average success index of PMv was actually higher than that of M1 and equally high than that of PMd (Kruskal-Wallis test, post-hoc comparisons p < 0.05, Supplementary Table 3). This effect was less pronounced in the Passive avatar tasks but even there, the average PMv-based decoder success index could exceed the one of M1 (e.g. in the Assisted Passive avatar task in monkey L.). Note that the PMv-based decoder success index showed a high variability over sessions.

**Figure 6:**
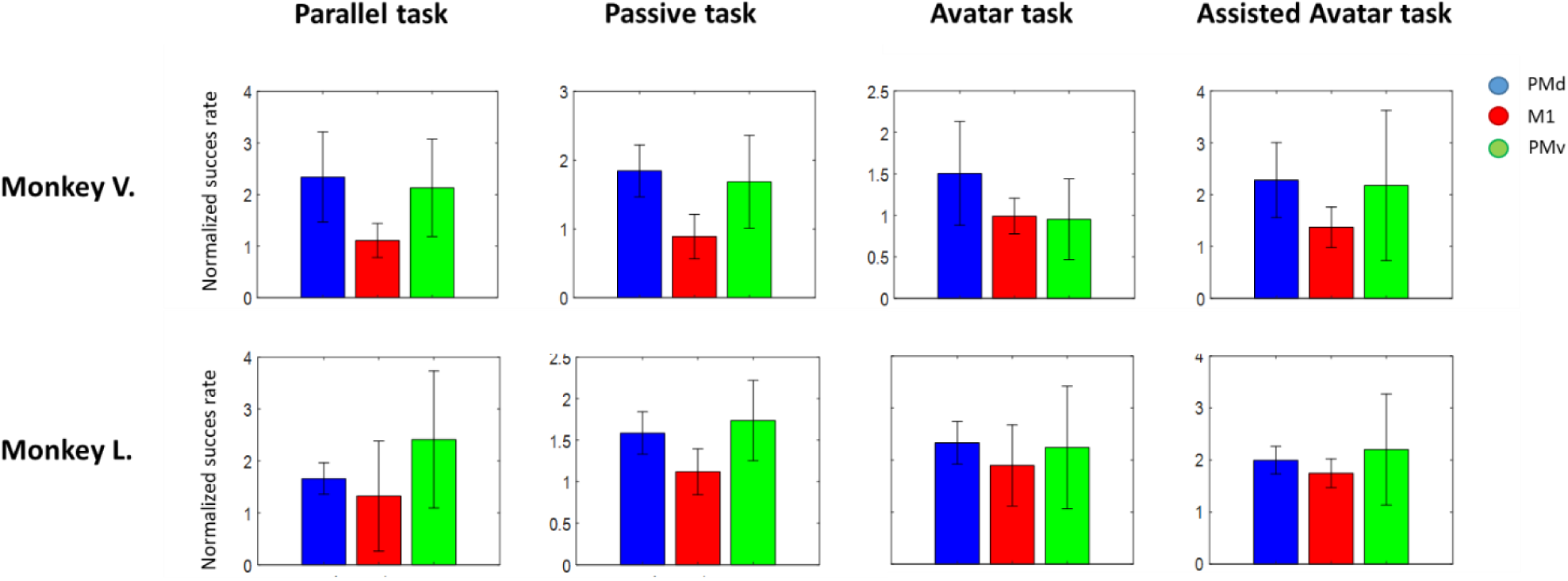
Success index of all tasks. For each monkey and each task, the success index is shown per area (blue = PMd, red = M1, and green = PMv) averaged over all recording sessions +/- SEM. The success index per session is the success rate of that session divided by the number of motion selective electrodes.

### Neural responses during the training phase and the online decoding phase

We also wanted to investigate to what extent the neural selectivity changed during online decoding compared to the training phase for the three areas tested. Figure 7 shows the average raw spiking activity recorded on three example electrodes during the training phase and the online decoding phase of the Parallel task. Note that in the online decoding phase, the monkeys were simultaneously performing the task while controlling the cursor movement. In the first example, there is a clear tuning for movement towards the targets on the right of the screen (d’ = 0.66) that was maintained in the online decoding phase (d’ = 0.68), although with a slightly lower spike rate (recorded in PMv, Fig. 7A). The activity of the second example electrode showed a similar neural tuning as the first example in the training phase (recorded in M1, Fig. 7B; d’ = 0.35). In the online decoding phase, the direction selectivity decreased but a preference for movements to the right was still present (d’ = 0.25), although less pronounced. In contrast, the last example showed less selectivity during the training phase (d’ = 0.33), but a high selectivity for the upper right targets during the online decoding phase (Recorded in M1, Fig. 7C; d’ = 0.46). Similar examples were found in all three areas in both monkeys.

**Figure 7:**
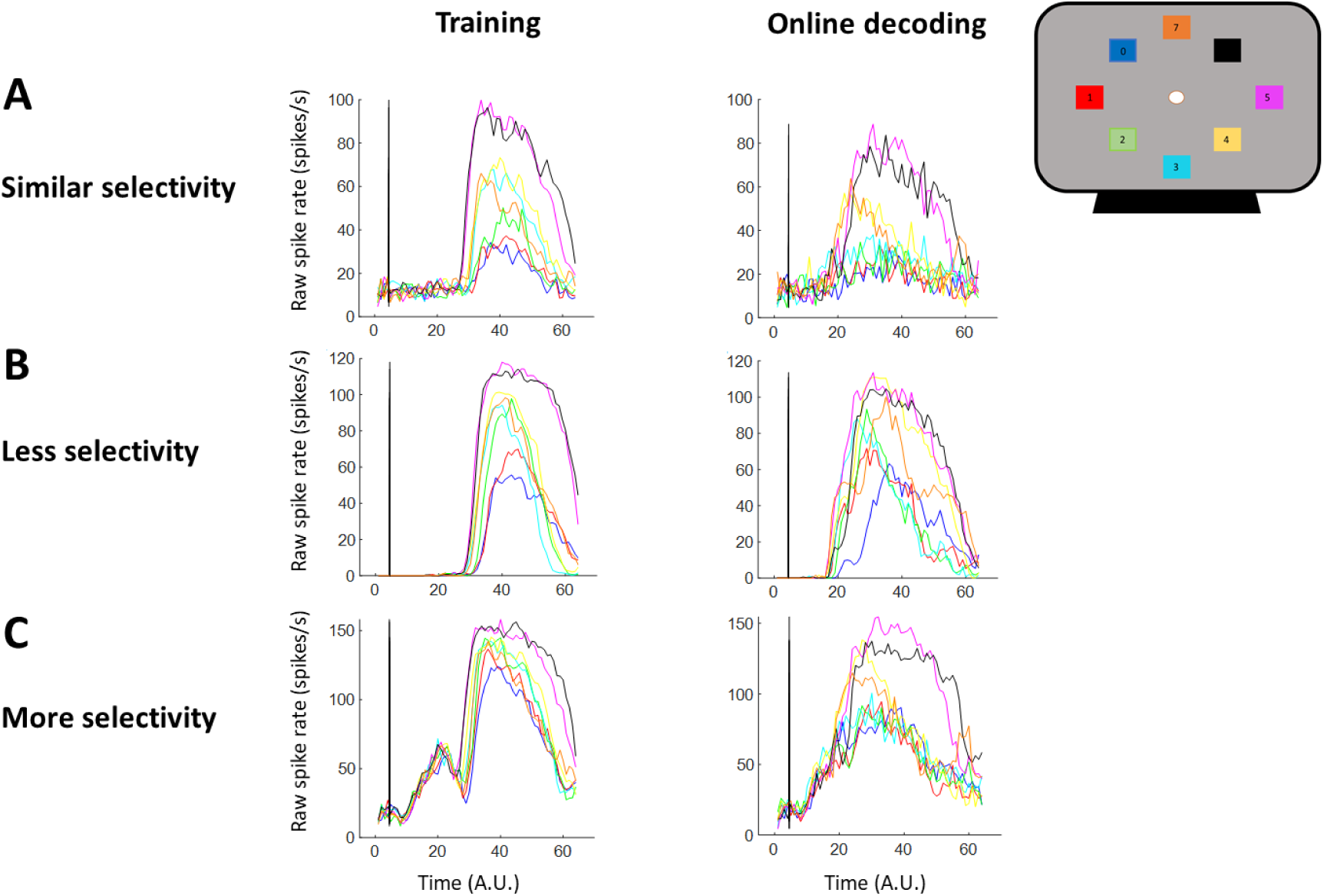
Example sites during training and online decoding. (A) Raw spike rate over time (in arbitrary units) of an example site in PMv during the training phase and the online decoding phase. Colors depict the different targets, as shown on the screen in the right upper corner. The vertical black line represents the onset of the cursor movement (B-C) Same as (A) but for two different example sites in M1.

To test whether PMv neurons alter their directional tuning more easily during online decoding compared to PMd and M1 neurons, we calculated the selectivity for direction of motion during the training phase in which the monkeys had no control over the cursor, and during the online decoding phase in which they actively controlled the cursor, as a d prime index (d’, see methods). We calculated the d’ of each site that was significantly modulated during both phases. In the Parallel task, we observed significant correlations between the d’ indices across sites during the training phase and the online decoding phase in both monkeys (r = 0.42 for PMv, r = 0.67 for PMd, and r = 0.61 for M1, all p<0.001 in monkey V., and r = 0.87 for PMv, r = 0.80 for PMd, and r = 0.88 for M1, all p<0.001 in monkey L.), indicating that most sites had a similar selectivity during passive observation of a cursor and active control of the cursor movement. Importantly, the monkeys executed the task in both phases, which may have contributed to the high correlations. Indeed, we observed much lower correlations when comparing the d’ indices during the training phase and the online decoding phase of the Passive cursor task (r = 0.22 for PMv, r = 0.25 for PMd, and r = 0.24 for M1, all p<0.05 in monkey V., and r = 0.12 for PMv, r = 0.21 for PMd, and r = 0.19 for M1, all p<0.05 in monkey L.).

To assess whether the neural activity changed its selectivity for direction of motion during online decoding compared to the training phase (i.e. whether neurons were recruited or disengaged during active cursor control), we calculated the change in d’ for each electrode with responsive activity in both phases. Overall, the majority of sites did not change their selectivity much when comparing both phases (d’ change values around zero, Supplementary Figure 1). Nonetheless, we found that more sites were recruited (i.e. showed an increased selectivity for direction of motion during the online decoding phase compared to the training phase, positive values in Supplementary Figure 1) during the passive cursor task compared to the parallel task. Note that the activity of these electrodes changed from either non-selective in the training phase to selective in the online decoding phase or selective in the training phase to even more selective in the online decoding phase. Next, we investigated whether PMv sites increased their selectivity more than PMd and M1 during online decoding. During both tasks, significantly larger changes in selectivity were observed in PMv compared to PMd (Supplementary Table 4). Notably, PMv and M1 sites were similarly recruited during online decoding in the passive cursor task in both monkeys. Thus, a subset of sites in each area showed an increase in the selectivity of motion direction during online decoding, especially during the passive cursor task, indicating that these neurons modulated their activity when the monkey started to actively control the cursor compared to passively observing the cursor movements.

### Population dynamics during training and decoding

To assess whether the neural population responded similarly during the training phase and the online decoding phase, we performed principal component (PC) analysis on the training data and then projected both the training and the online decoding data onto the training PCs. Figure 8A shows the evolution of the population responses during the training phase (light shade) and the online decoding phase (dark shade) in a shared neural space for an example recording session during each task. For each area, we observed highly similar representations in a three-dimensional subspace, suggesting alignment between the training and the online decoding phase. To quantify the alignment between the training and the online decoding activity, we computed an alignment index (AI) for each area of each recording session with the subspace overlap analysis (Elsayed et al. 2016). The AI was calculated by quantifying the training data variance captured by the first ten decoding PCs, normalized by the training data variance captured by the first ten training PCs (see methods). The index ranges from 0 (perfectly orthogonal subspaces) to 1 (perfectly aligned subspaces). For both monkeys, we found similar high indices in both tasks and in all three areas (Figure 8B), indicating that the population responses during the training phase and the online decoding phase exhibited remarkably similar dynamics, and thus relied on largely overlapping neural population representations.

**Figure 8:**
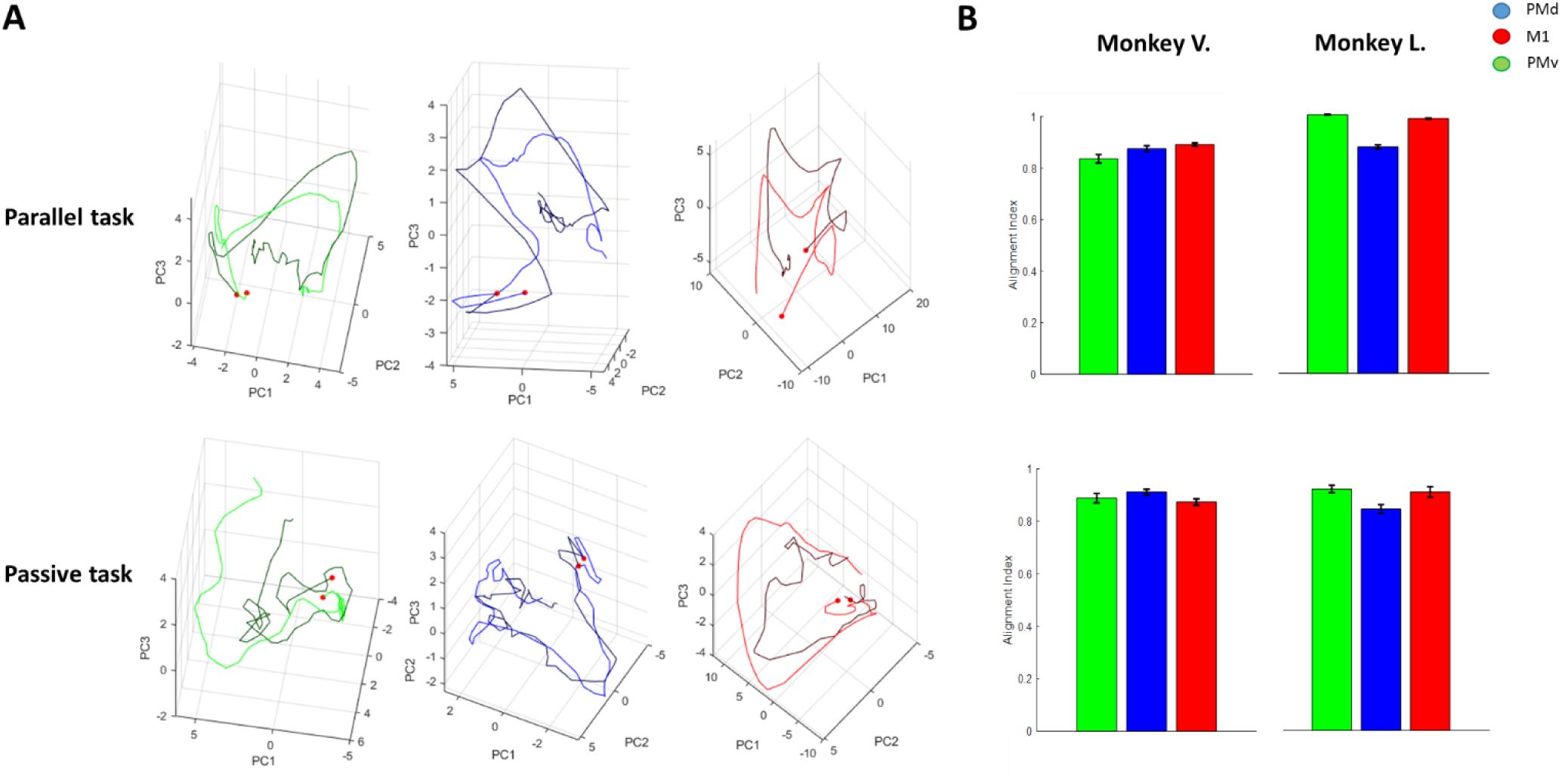
Population dynamics during the training phase and the online decoding phase. (A) Neural trajectories during the training phase (light shade) and the online decoding phase (dark shade) for an example recording session in Monkey V. during the Parallel and the Passive cursor task. The red dot indicates the first time point of the trajectory, i.e. the start of the cursor movement. (B) Average alignment index (+/- SEM) of the neural subspaces during the training phase and the online decoding phase (amount of sessions per area and per monkey range from 11 to 38). The different colors indicate the areas (blue = PMd, red = M1, and green = PMv).

## Discussion

No study has investigated the potential of F5c, a subsector of PMv, for online 2D cursor control. Therefore, we systematically compared the efficacy of M1, PMd and F5c using chronically implanted multi-electrode arrays in three different tasks in macaque monkeys. Although the overall performance of the BMI heavily depended on the number of motion selective electrodes in each area, decoders based on F5c activity reached a similar performance level as decoders based on PMd activity, and could even achieve a higher performance per electrode compared to M1-based decoders. For robot avatar control, we obtained lower success rates than for 2D cursor control. In all three areas, the neuronal tuning for direction of motion and the population dynamics were highly similar between the training phase and the online decoding phase. Overall, our findings demonstrate the usefulness of an invasive BMI based on F5c activity for online cursor and assistive robot avatar control.

It may appear surprising that an area such as F5c, which has been associated with high-level motor planning of grasping movements and action recognition, can be used efficiently for controlling a cursor moving in eight possible directions on a display. However, in our previous study (De Schrijver, Decramer, and Janssen 2024) we noticed that the average response of F5c neurons in a delayed grasping task mimicked that of M1, with relatively weak responses after object presentation and in the delay period, but a strong increase in activity after the lift of the hand. Furthermore, 62% of all task-modulated F5c neurons also responded during passive fixation of action videos (Action Observation/Execution neurons or AOENs). The large majority of these AOENs (79%) were tuned to a specific epoch of the action video and 74% also responded to a simple ellipse moving in the visual field – even in the absence of a graspable object. In the light of these findings, it is reasonable that in our Passive cursor task, a simple cursor moving on a display was sufficient to recruit a large population of F5c neurons, the large majority of those being AOENs. AOENs have also been described in PMd and M1 (Papadourakis and Raos 2019; Tkach, Reimer, and Hatsopoulos 2007), and preliminary data suggest that simple stimuli can also activate AOENs in these other cortical areas. Note that previous studies have decoded different grasp types (precision compared to power grip) from PMv activity (Townsend, Subasi, and Scherberger 2011), which was most likely primarily F5p, the subsector of F5 that is most heavily and directed connected with M1 and the spinal cord (Borra et al. 2010).

We reasoned that the F5c decoding performance might be improved by using a robot avatar instead of a simple cursor, since the mirror system in humans is activated by both robot arm and hand grasping objects (Gazzola et al. 2007). Contrary to our expectations, but in line with (Nelissen et al. 2005), the decoding performance in the Passive avatar task did not exceed that in the Passive cursor task in any of the three areas, despite the more realistic effector used in this task. It is likely that the absence of realistic depth cues (such as binocular disparity) in the Passive avatar task was partially responsible for the lack of decoding performance improvement.

Performance in any of the tasks we employed depended on a number of external factors of which the number of motion selective electrodes for each recording session might have been the most important. Over the course of several months of experiments, we observed large changes in the number of electrodes with spiking activity, predominantly in F5c in both monkeys (and in PMd in monkey V.). The clear improvement in F5c-based decoder performance over time in the Passive and Parallel tasks in both monkeys can most likely be explained by a marked increase in the number of motion selective electrodes. In addition, the initial decoding sessions occurred not long after array implantation, which may have been responsible for a lack of stability in the recording signal. Thus, although the stronger increase in decoder performance in F5c compared to the other two areas may suggest that the monkeys needed more time to learn cursor control with their F5c activity, other factors largely explain the change in online decoding performance. When we controlled for the number of electrodes with selective spiking activity that correlated with the cursor movements in the training phase, we unexpectedly even measured a superior decoder performance based on F5c activity over M1 activity in most tasks. In other words, per electrode with selective spiking activity, decoders using F5c activity performed better in 2D cursor control than decoders using M1 activity.

When addressing the performance of any decoding model, the chosen set of hyperparameters influences critical aspects of the pipeline. In our case, these hyperparameters influenced the number of selected channels through the kinematic threshold, the descriptive power of the model through the choice of the number of behaviorally relevant states and how robust the system is to overfitting through the regularization parameters. We fixed the hyperparameters for all recording sessions, regardless of monkey, area, and task, to enable fair comparisons of the performance across different brain areas. Optimizing the hyperparameters per recording session would likely have led to better performances. Additionally, the arrays were not implanted in the exact same location in the subsector of each area in the different monkeys, which also may have contributed to the observed differences in performance between the two monkeys.

A small but noticeable number of sites modulated its activity when switching from passively observing the cursor to actively controlling the cursor movements, emphasizing the importance of retraining the decoder during online decoding. Sites that lost their selectivity for direction of motion during online decoding could be discarded, while sites with increased selectivity could be included in the recalculation of the decoder. In all three areas, the majority of modulated sites increased their selectivity for the direction of motion during active BMI control. Importantly, PMv and M1 neurons increased their selectivity more than PMd during the passive cursor task, indicating that PMv and M1 show a higher adaptability to the motor BMI. Note, however, that the majority of sites responded similarly during training and online decoding. Indeed, we found a high overlap between training and decoding subspaces in each area, indicating largely overlapping neural population representations during training and decoding.

We did not observe any evidence for more intuitive cursor control or faster learning based on F5c activity compared to PMd or M1 activity. In fact, F5c-based decoder performance in the Passive cursor task improved remarkably over time in both monkeys while PMd- and M1-based decoder performance appeared more stable. However, this finding is confounded by the change in the signal quality on the F5c arrays. Over the course of several weeks, the number of channels with spiking activity on the F5c arrays increased significantly and thereby also improved the performance.

Consistent with previous studies (Carmena et al. 2003), we also noticed that reliable cursor control requires at least 30-40 motion selective electrodes. Future studies could combine the neural activity of all three areas to train the decoder, hence increasing the amount of input for the decoder. Additionally, using the same input on a daily basis will induce a steeper learning curve since alternating between inputs (i.e. changing the brain area that is used for decoding daily) might disrupt learning performance (Moritz and Fetz 2011). Moreover, integrating information from multiple brain areas could significantly improve decoder performance (Gallego, Makin, and McDougle 2022) and enable the exploration of the neural interplay between these areas during online BMI control. Finally, more complex and realistic experimental tasks (in real-life or in virtual reality) are needed to advance towards clinical applications such as fast and flexible wheelchair control.

## Acknowledgments

This work was supported by Fonds Wetenschappelijk onderzoek (FWO) grant G.097422N and KU Leuven grants C14/18/100 and C14/22/134. We thank Stijn Verstraeten, Marc De Paep, Wouter Depuydt, Inez Puttemans, and Christophe Ulens for technical assistance. We thank Astrid Hermans and Sara De Pril for administrative support.

## Data availability

Processed data will be made available on the Dryad server.

