## Supplementary material for "An intracortical brain-machine interface based on macaque ventral premotor activity"

| Pipeline step | Parameter | Description | Value Range | Final value |
|---|---|---|---|---|
| Kinematic Filter | $\gamma$ | Threshold used for kinematic selectivity filtering | 0.3-0.75 | 0.3 |
| PSID | n1 | Number of behaviorally-relevant hidden states | 2 - 10 | 10 |
| PSID | nx | Total number of hidden states | 10 | 10 |
| PSID | i | The subspace horizon used for modeling | 5 | 5 |
| Nymstrom Kernel | $\delta$ | Regularization parameter for the Nyström RBF kernel | 0.005 - 1.0 | 0.341 |
| Nymstrom Kernel | n_components | Number of components used in the Nyström approximation | 700 | 700 |
| Nymstrom Kernel | random_state | Random seed for the Nyström approximation | 42 | 42 |
| Ridge Regression | L2 | Regularization strength for the ridge regression model | 0.1 | 0.1 |

Supplementary Table 1: Hyperparameters of the decoder.

| | Monkey V. | | | Monkey L. | | |
|---|---|---|---|---|---|---|
| | PMv | PMd | M1 | PMv | PMd | M1 |
| Parallel Task | 13 | 16 | 13 | 13 | 17 | 18 |
| Passive cursor Task | 11 | 10 | 12 | 14 | 14 | 11 |
| Passive avatar Task | 9 | 7 | 10 | 7 | 7 | 7 |
| Passive assisted Avatar Task | 14 | 12 | 13 | 7 | 7 | 7 |

Supplementary Table 2: Number of recording sessions per area, task and monkey, where the listed area indicates the one used for decoder training and online decoding.

| Kruskal-Wallis test on normalized success rate (and post-hoc tests) | Monkey V. | | | | Monkey L. | | | |
|---|---|---|---|---|---|---|---|---|
| | PMv – M1 | PMv - PMd | PMd – M1 | All Areas | PMv – M1 | PMv - PMd | PMd – M1 | All Areas |
| Parallel Task | p = 0.0126 | p = 0.9496 | p = 0.0015 | H(2) = 13.62<br>p = 0.0011 | p = 4.385e-04 | p = 0.7262 | p = 0.0064 | H(2) = 16.73<br>p = 2.3286e-04 |
| Passive Task | p = 0.0060 | p = 0.8254 | p = 0.0004 | H(2) = 16.63<br>p = 2.4501e-04 | p = 0.0012 | p = 0.9282 | p = 0.0075 | H(2) = 14.09<br>p = 8.7073e-04 |
| Avatar Task | p = 0.9271 | p = 0.1264 | p = 0.3232 | H(2) = 4.28<br>p = 0.1176 | p = 0.8679 | p = 0.9630 | p = 0.5993 | H(2) = 1.28<br>p = 0.5282 |
| Assisted Avatar Task | p = 0.1233 | p = 0.6047 | p = 0.0071 | H(2) = 9.59<br>p = 0.0083 | p = 0.1360 | p = 0.8460 | p = 0.5100 | H(2) = 4.01<br>p = 0.1343 |

Supplementary Table 3: Statistics of the success index of all tasks for each monkey

| Kolmogorov-Smirnov test on the distributions of the d' change between training and online decoding | | Monkey V. | | | Monkey L. | | |
|---|---|---|---|---|---|---|---|
| | | PMv – M1 | PMd – PMv | M1 – PMd | PMv – M1 | PMd – PMv | M1 – PMd |
| Parallel Task | KS statistics | D = 0.1774<br>p = $9.4901e^{-16}$ | D = 0.1047<br>p = $6.5067e^{-04}$ | D = 0.0825<br>p = 0.0013 | D = 0.0759<br>p = $4.3203e^{-04}$ | D = 0.1827<br>p = $8.2709e^{-21}$ | D = 0.1783<br>p = $3.5792e^{-28}$ |
| Passive cursor Task | KS statistics | D = 0.0589<br>p = 0.2113 | D = 0.0661<br>p = 0.2629 | D = 0.1091<br>p = 0.0010 | D = 0.0512<br>p = 0.8213 | D = 0.2364<br>p = $3.6410e^{-10}$ | D = 0.2722<br>p = $3.0313e^{-15}$ |

Supplementary Table 4: Statistics of the distributions of the d' changes in the training phase and the online decoding phase to assess the change in selectivity for direction of motion in the Parallel and Passive task.

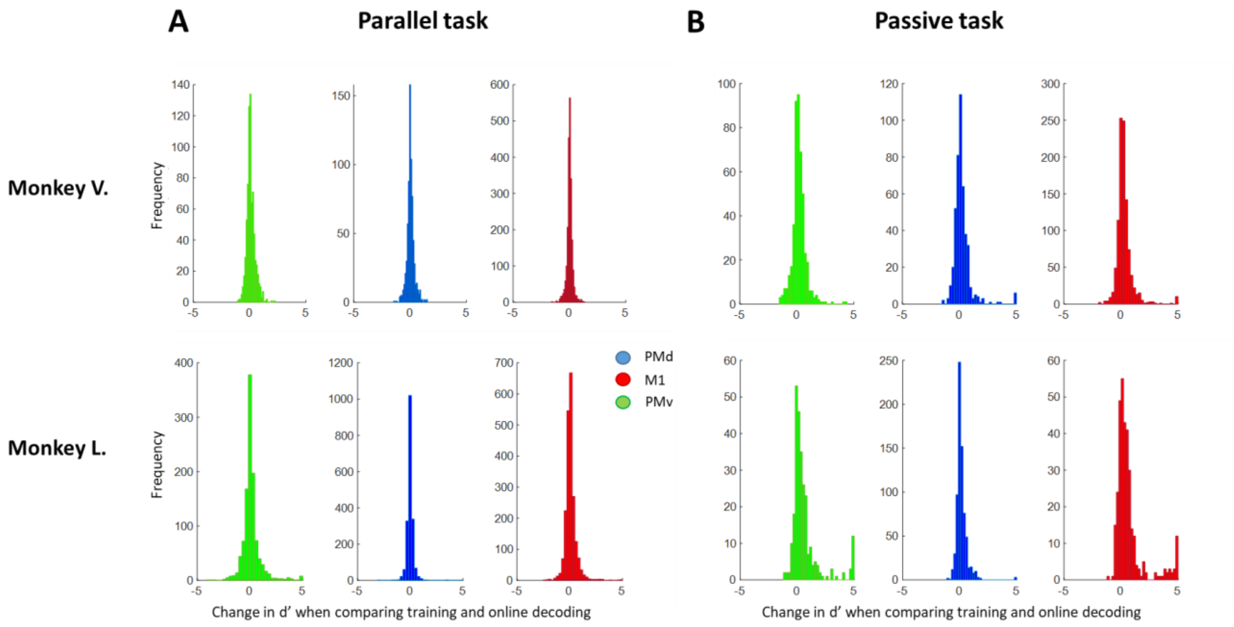


Supplementary Figure 1: Difference in selectivity for motion direction during training and online decoding. Change in d' for all responsive sites when comparing the training phase and the online decoding phase during the Parallel task (A) and the Passive task (B). A positive change in d' indicates more selectivity during online decoding, whereas a negative change indicates more selectivity during training. The colors represent the three areas (green = PMv, blue = PMd, and red = M1). For visualization purposes, a cut-off was made at -5 and 5.
